# Curcumin micronization by supercritical fluid: *in vitro* and *in vivo* biological relevance

**DOI:** 10.1101/2021.07.08.451641

**Authors:** Adrieli Sachett, Matheus Gallas-Lopes, Radharani Benvenutti, Matheus Marcon, Gean Pablo S. Aguiar, Ana Paula Herrmann, J. Vladimir Oliveira, Anna M. Siebel, Angelo Piato

**Affiliations:** Programa de Pós-graduação em Neurociências, Instituto de Ciências Básicas da Saúde, Universidade Federal do Rio Grande do Sul (UFRGS), Porto Alegre, RS, Brazil; Departamento de Farmacologia, Instituto de Ciências Básicas da Saúde, Universidade Federal do Rio Grande do Sul (UFRGS), Porto Alegre, RS, Brazil; Programa de Pós-graduação em Farmacologia e Terapêutica, Instituto de Ciências Básicas da Saúde, Universidade Federal do Rio Grande do Sul (UFRGS), Porto Alegre, RS, Brazil; Programa de Pós-Graduação em Ciências Ambientais, Universidade Comunitária da Região de Chapecó (Unochapecó), Chapecó, SC, Brazil; Departamento de Engenharia Química e de Alimentos, Universidade Federal de Santa Catarina (UFSC), Florianópolis, SC, Brazil

**Keywords:** supercritical fluids, unpredictable chronic stress, oxidative damage, antioxidant, curcumin, micronization, zebrafish

## Abstract

Curcumin, a polyphenol extracted from the rhizome of *Curcuma longa* L. (Zingiberaceae), is shown to have antioxidant, anti-inflammatory, neuroprotective, anxiolytic, and antidepressant properties in both preclinical and clinical studies. However, its low bioavailability is a limitation for its potential adoption as a therapeutic agent. The process of micronization can overcome this barrier by reducing the particle size and increasing the dissolution rate, potentially improving the bioavailability of the compounds of interest. In this study, we compared the *in vitro* antioxidant effects of curcumin (CUR) and micronized curcumin (MC) and studied their effects on behavioral and neurochemical parameters in zebrafish submitted to unpredictable chronic stress (UCS). MC (1 g/L) presented higher antioxidant activity *in vitro* as compared to CUR, as measured by iron-reducing antioxidant power (FRAP), 1,1-diphenyl-2-2-picyryl-hydrazyl radical removal (DPPH), and deoxyribose tests. UCS increased total distance traveled in the social interaction test (SI), while decreased crossings, time, and entries to the top area in the novel tank test (NTT). No effects of UCS were observed in the open tank test (OTT). The behavioral effects induced by UCS were not blocked by any curcumin preparation. UCS also decreased non-protein thiols (NPSH) levels, while increased glutathione reductase (GR) activity and thiobarbituric acid reactive substances (TBARS) levels on zebrafish brain. MC presented superior antioxidant properties than CUR *in vivo*, blocking the stress-induced neurochemical effects. Although this study did not measure the concentration of curcumin on the zebrafish brain, our results suggest that micronization increases the bioavailability of curcumin, potentiating its antioxidant activity both *in vitro* and *in vivo*. Our study also demonstrates that counteracting the oxidative imbalance induced by UCS is not sufficient to block its behavioral effects.

## 1. Introduction

Stress is an adaptative process by which the body reacts to an external stimulus or threat. Physiological, behavioral, and metabolic adaptations through the activation of the sympathetic autonomic nervous system and the hypothalamic-pituitary-adrenal axis (HPA) are triggered so that an organism adequately responds to that stimulus (Kaufmann et al., 2016; Mcewen, 2006; McEwen et al., 2015). However, when the individual is exposed to chronic stress, depletion of the adaptive response can occur with deleterious results. In this case, hyperactivation of the sympathetic autonomic nervous system and HPA axis, increased cortisol levels, dysfunction of neurotransmitter systems, neuroinflammation, defects in neurogenesis and synaptic plasticity, mitochondrial dysfunction, redox state imbalance, and oxidative damage may be present. These complex neurobiological changes can predispose the individual to mental disorders such as anxiety and depression (Cernackova et al., 2020; Popoli et al., 2012). The neurobiological basis of mental disorders is not fully understood, but studies have already shown that oxidative status imbalances are present in animal models and humans (Avery, 2011; Fedoce et al., 2018; Harwell, 2007; Morris et al., 2020; Picard et al., 2018; Valko et al., 2007). Therefore, compounds with antioxidant activity are potential candidates, as a complement to the non-pharmacological and pharmacological therapeutic approaches.

Curcumin is one of the compounds extracted from *Curcuma longa* L. (Zingiberaceae) roots, widely used in Asian countries as a food coloring and seasoning component. Several works have demonstrated the potent antioxidant activity of this compound, both *in vitro* and *in vivo* (Ak and Gülçin, 2008; Menon and Sudheer, 2007). Curcumin can attenuate intracellular production of reactive oxygen species (ROS), increase antioxidant enzyme activity, and protect mitochondria from oxidative damage in rats (Wei et al., 2006; Zhu et al., 2016). Moreover, this compound has shown anti-inflammatory, neuroprotective, and immunomodulatory effects (Bhutani et al., 2009; Kulkarni et al., 2008; Reeta et al., 2010; Wang et al., 2008, 2014; Yadav et al., 2005; Zhao et al., 2014).

However, curcumin has low bioavailability due to poor absorption, rapid metabolism, and quick systemic elimination, which compromise its therapeutic use for neuropsychiatric disorders (Anand et al., 2007; Yang et al., 2005). The micronization process reduces the size of particles modifying the conformation of crystal structure, changing the physical structure, and enhancing solubility and dissolution rates. As expected, these effects could lead to an improvement in the bioavailability of curcumin.

Previous studies have shown that micronization reduced the particle size of *Panax notoginseng* saponins, changing dissolution/release and increasing concentration in the plasma of rats compared with the larger particle size preparations (Liang et al., 2021). Also, the micronized purified flavonoid fraction (diosmin + hesperidin) showed a significant anticoagulant effect, increasing platelet disaggregation in rats (McGregor et al., 1999). In humans, micronized resveratrol (SRT501) showed higher plasmatic concentration, providing measurable resveratrol levels in liver tissue of patients with colorectal cancer and hepatic metastases. Moreover, STR501 increased the cleaved caspase-3 (an apoptosis biomarker) in malignant hepatic tissue (Howells et al., 2011). Our group has shown that micronization decreased the minimum effective concentration of N-acetylcysteine (NAC) required to exert an anxiolytic effect in zebrafish (Aguiar et al., 2017). Also, micronized curcumin and resveratrol, but not non-micronized compounds, showed similar effects as the antiepileptic drugs diazepam and valproate, reducing seizure occurrence and slowing seizure progression in the PTZ-induced seizure model in zebrafish (Almeida et al., 2021; Bertoncello et al., 2018; Decui et al., 2020).

Considering the role of chronic stress in neuropsychiatric disorders and the potential neuromodulatory effects of curcumin, this study aimed to compare the effects of curcumin and micronized curcumin on behavioral and neurochemical parameters in adult zebrafish submitted to the unpredictable chronic stress. In addition, we compared the antioxidant effects of both preparations *in vitro*.

## 2. Materials and methods

### 2.1 Drugs

Curcumin was obtained from Sigma-Aldrich® (CAS 458-37-7) (St. Louis, MO, USA), and its micronization was carried out at the Laboratory of Thermodynamics and Supercritical Technology (LATESC) of the Department of Chemical and Food Engineering (EQA) at UFSC. Both curcumins preparations were dissolved in 1% DMSO (Dimethyl sulfoxide anhydrous) obtained from Sigma-Aldrich® (CAS 67-68-5) and diluted in injection water (Samtec biotecnologia®, SP, Brazil) acquired from a commercial supplier. Other reagents for neurochemical analysis were obtained from Sigma-Aldrich®.

### 2.2 Curcumin micronization with the solution enhanced dispersion by supercritical fluids (SEDS)

The SEDS experimental equipment and procedure used to micronize curcumin with supercritical carbon dioxide (CO_2_) as an anti-solvent was described in detail in previous studies (Dal Magro et al., 2017; Machado et al., 2014). A schematic diagram of the experimental apparatus is presented in Figure 1. The process parameters adopted in the present report were based on previous data: solute concentration of 20 mg/mL, temperature at 35 °C, anti-solvent flow rate of 20 mL/min, solution flow rate of 1 mL·min ^-1^, and operating pressure of 8 MPa (Aguiar et al., 2018, 2017, 2016; Bertoncello et al., 2018).

**Fig. 1.**
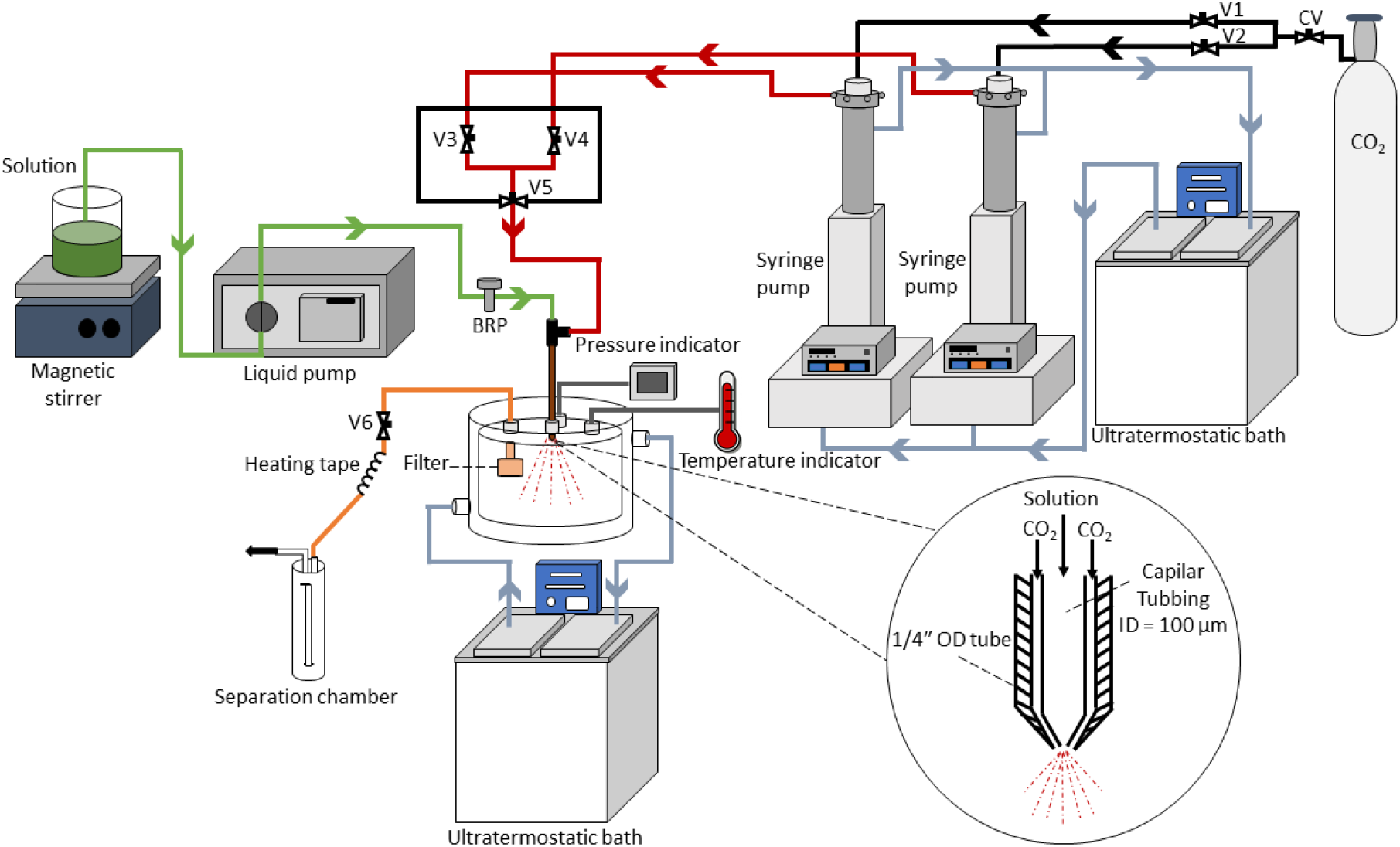
Schematic diagram of the experimental apparatus using the solution enhanced dispersion by supercritical fluids technique (SEDS). CV (check valve), V1, V2, V3, and V4 (ball valve), V5 and V6 (needle valve), and BRP (backpressure regulator).

### 2.3 Morphology and determination of particle size

CUR and MC samples were submitted to morphological analysis by Scanning Electron Microscopy (SEM) (JEOL JSM-6390LV United States), with 10 kV power and 300-500 objectives, to determine particle morphology and Meter Size software (version 1.1) was used to determine the mean particle size (Aguiar et al., 2016; Bertoncello et al., 2018).

### 2.4 Differential scanning calorimetry

The melting point of the CUR and MC was determined using a system of differential scanning calorimetry (DSC) (Jade-DSC, Perkin Elmer). The samples (5–10 mg) were prepared in an aluminum pan, and DSC measurements were performed by heating at 30 to 200 °C at a rate of 10 °C/min in an inert atmosphere (N2 flow: 10 mL/min) (Aguiar et al., 2016; Bertoncello et al., 2018).

### 2.5 *In vitro* antioxidant activity

To analyze the antioxidant activity *in vitro*, the following experimental groups were tested: 1% DMSO; ascorbic acid (0.0625, 0.25 and 1 g/L, as positive control); CUR (0.0625, 0.25 and 1 g/L) and MC (0.0625, 0.25 and 1 g/L). These concentrations were based on previous studies (Bertoncello et al., 2018; Gilhotra and Dhingra, 2010; Xu et al., 2005). All analyses were performed in duplicate with an n=5. A schematic diagram of the experimental tests is presented in Figure 2.

**Fig. 2.**
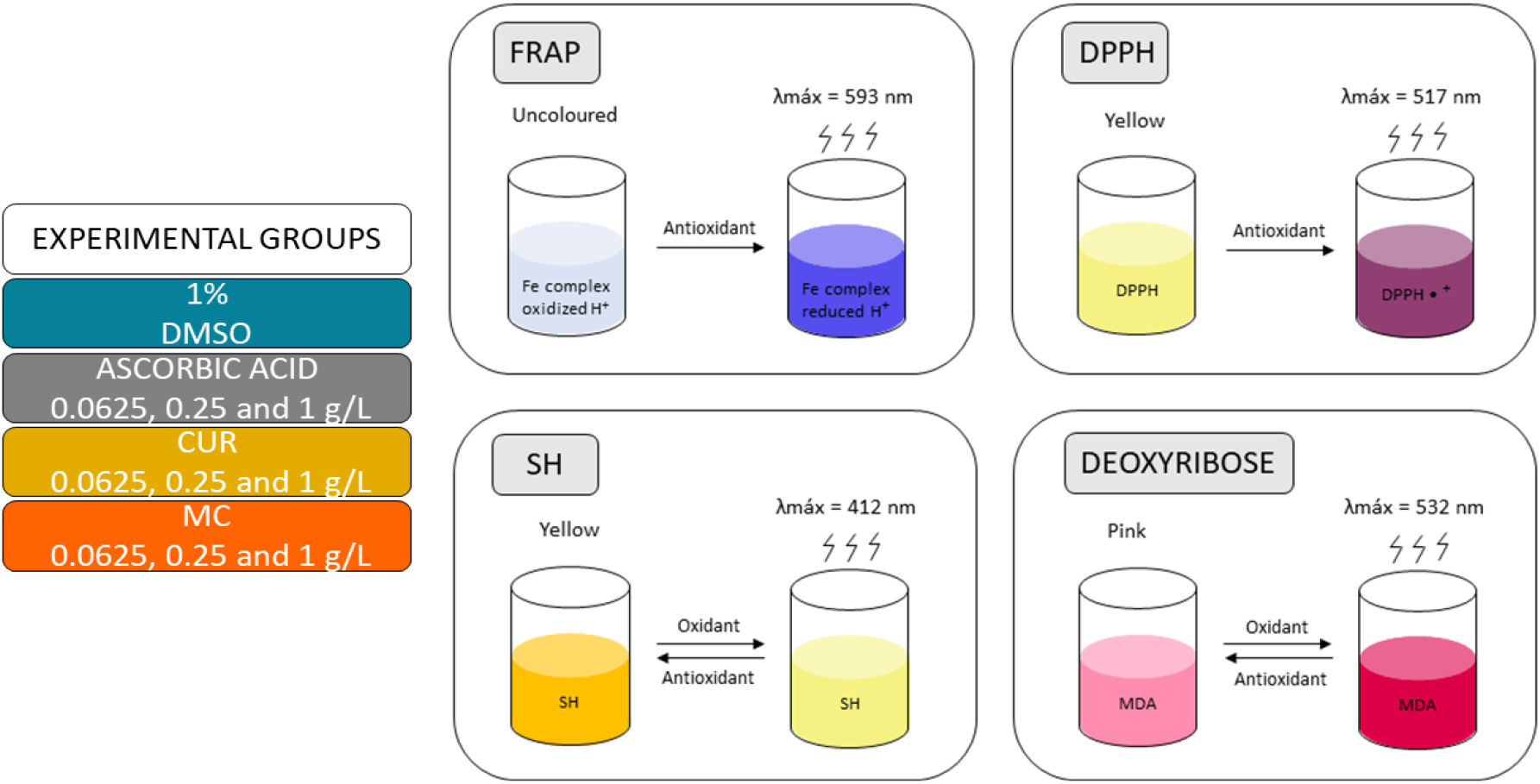
Experimental design of the *in vitro* assays: FRAP (iron-reducing antioxidant power), DPPH (1,1-diphenyl-2-2-picyryl-hydrazyl radical removal), GSH (protection against glutathione oxidation), deoxyribose assay. DMSO (dimethyl sulfoxide), CUR (curcumin) and MC (micronized curcumin).

#### 2.5.1 Determination of iron-reducing antioxidant power (FRAP)

The antioxidant power was assessed by the reduction of Fe^3+^ to Fe^2+^. The samples were incubated at 37 °C for 15 min with a reaction medium containing: 10 mM 2,4,6-Tripyridyl-Triazine (TPTZ) + 40 mM hydrochloric acid (HCl), FeCl_3_. 20 mM 6H_2_O and 300 mM acetate buffer (10:1:1 ratio). The absorbance was read at 593 nm. Blanks for each sample were incubated without TPTZ (Benzie and Strain, 1996). The detailed protocol is available at protocols.io (Sachett et al., 2021a).

#### 2.5.2 1,1-diphenyl-2-2-picyryl-hydrazyl radical removal test (DPPH)

The scavenging capacity of free radicals was assessed by the DPPH assay. The samples at different concentrations were incubated for 24 h in the dark with methanol and DPPH (0.24 mg/mL). DPPH and methanol, without samples, were used as controls. Methanol was used as a blank. After incubation, the absorbance was read at 517 nm. The effective concentration 50 (EC50) for each extract, which expresses the minimum amount of the extract capable of reducing the initial concentration of DPPH radical by 50%, was calculated by non-linear regression using GraphPad Prism version 8 (Brand-Williams et al., 1995). The detailed protocol is available at protocols.io (Sachett et al., 2021b).

#### 2.5.3 Protection against glutathione oxidation (GSH)

To quantify the presence of sulfhydryl groups after oxidation induced by hydrogen peroxide (H_2_O_2_). Each sample was incubated for 30 min in the dark, with a reaction medium containing potassium phosphate buffer (TFK) (200 mM, pH 6.4) and H_2_O_2_ (5 mM). Afterward, the mixture was added to 5,5’-dithiobis-(2-nitrobenzoic acid) (DTNB) (10 mM) and the absorbance was read at 412 nm after 5 min. Samples without GSH were used as sample blank and the incubation medium without sample was used as a control (Ellman, 1959). The detailed protocol is available at protocols.io (Sachett et al., 2021c).

#### 2.5.4 Deoxyribose assay

The production of malondialdehyde (MDA) after oxidation of deoxyribose by hydroxyl radical (OH·) was measured through the TBARS. The samples were incubated at 37°C for 1 h, with a reaction medium containing: KH_2_PO_4_-KOH (50 mM, pH 7.4), deoxyribose (60 mM), FeCl_3_ (1 mM), ethylenediaminetetraacetic acid (EDTA) (1.04 mM), ascorbic acid (2 mM), and H_2_O_2_ (10 mM). Then, 1% thiobarbituric acid (TBA) and 25% HCl were added to the mix and heated in a water bath at 100 °C for 15 min. The absorbance was read at 532 nm. Samples without deoxyribose were used as sample blank and the incubation medium without sample was used as a control (Halliwell et al., 1987). The detailed protocol is available at protocols.io (Sachett et al., 2021d).

### 2.6. Animals

All procedures were approved by the institutional animal welfare and ethical review committee at the Federal University of Rio Grande do Sul (UFRGS) (approval #35279/2018). The animal experiments are reported in compliance with the ARRIVE guidelines 2.0 (Percie du Sert et al., 2020). Experiments were performed using 108 male and female (50:50 ratio) short-fin wild-type zebrafish, 6 months old, weighing 300 to 400 mg. Adult animals were obtained from the colony established at the Biochemistry Department of UFRGS and maintained in our animal facility (Altamar, SP, Brazil) in a light/dark cycle of 14/10 hours for at least 15 days before tests. Fish were transferred to 16-L home tanks (40 x 20 x 24 cm) with non-chlorinated water kept under constant mechanical, biological, and chemical filtration at a maximum density of two animals per liter. Tank water satisfied the controlled conditions required for the species (26 ± 2 °C; pH 7.0 ± 0.3; dissolved oxygen at 7.0 ± 0.4 mg/L; total ammonia at <0.01 mg/L; total hardness at 5.8 mg/L; alkalinity at 22 mg/L CaCO_3_; and conductivity of 1500–1600 μS/cm). Food was provided twice a day (commercial flake food (Poytara®, Brazil) plus the brine shrimp *Artemia salina*).

The animals were allocated to the experimental groups following block randomization procedures to counterbalance the sex, the two different home tanks, and the test arenas between the groups. Each experimental group was originated from two identical home tanks. Animal behavior was video recorded and analyzed with the ANY-Maze tracking software (Stoelting Co., Wood Dale, IL, USA) by researchers blinded to the experimental groups. All tests were performed between 08:00 and 12:00 a.m. The sex of the animals was confirmed after euthanasia by dissecting and analyzing the gonads. For all experiments, no tank and sex effects were observed, so data were pooled together.

After the tests, animals were euthanized by hypothermic shock according to the AVMA Guidelines for the Euthanasia of Animals (Leary and Johnson, 2020). Briefly, animals were exposed to chilled water at a temperature between 2 and 4 °C for at least 2 min after loss of orientation and cessation of opercular movements, followed by decapitation as a second step to ensure death.

### 2.7 Drug administration

Intraperitoneal (i.p.) injections were applied using a Hamilton Microliter™ Syringe (701N 10 μL SYR 26s/2”/2) x Epidural catheter 0.45 x 0.85 mm (Perifix®-Katheter, Braun, Germany) x Gingival Needle 30G/0.3 x 21 mm (GN injecta, SP, Brazil). The injection volume was 1 μL/100 mg of animal weight. The animals were anesthetized by immersion in a solution of tricaine (300 mg/L, CAS number 886-86-2) until loss of motor coordination and reduction of respiratory rate. The anesthetized fish were gently placed in a sponge soaked in water placed inside a petri dish, with the abdomen facing up and the fish’s head positioned on the sponge’s hinge. The needle was inserted parallel to the spine in the abdomen’s midline posterior to the pectoral fins. This procedure was conducted in approximately 10 seconds. The behavioral tests took place 24 hours after the last injection. Drug solutions were prepared daily.

### 2.8 Unpredictable chronic stress (UCS)

UCS was carried out based on previous studies (Bertelli et al., 2021; Marcon et al., 2019; Mocelin et al., 2019; Piato et al., 2011). The experimental design is presented in Figure 3 and the schedule and stressors are detailed in the supplementary material (Table S1). Initially, fish were divided into control (non-stressed group, S-) and UCS (stressed group, S+). After seven days, the experimental groups were subdivided into DMSO (1% DMSO), CUR (10 mg/kg), and MC (10 mg/kg) (this dose was chosen based on the concentration with the best antioxidant effect *in vitro*). The animals were anesthetized daily and injected at 2:00 p.m. (as described above) and then returned to the home tanks. The animals’ weight was checked on the 1^st^, 7^th^, and 14^th^ day and an average between the weights of each tank was used to calculate the injection volume. After the UCS, fish were submitted to the SI, NTT, and OTT, performed on the 15^th^, 16^th^, and 17^th^ days, respectively. On the 15^th^ and 16^th^ days, after the behavioral tests, the animals were also injected with the corresponding treatments. On the 17^th^ day, immediately after the OTT, fish were euthanized, and the brain was dissected and homogenized for the biochemical assays.

**Fig. 3.**
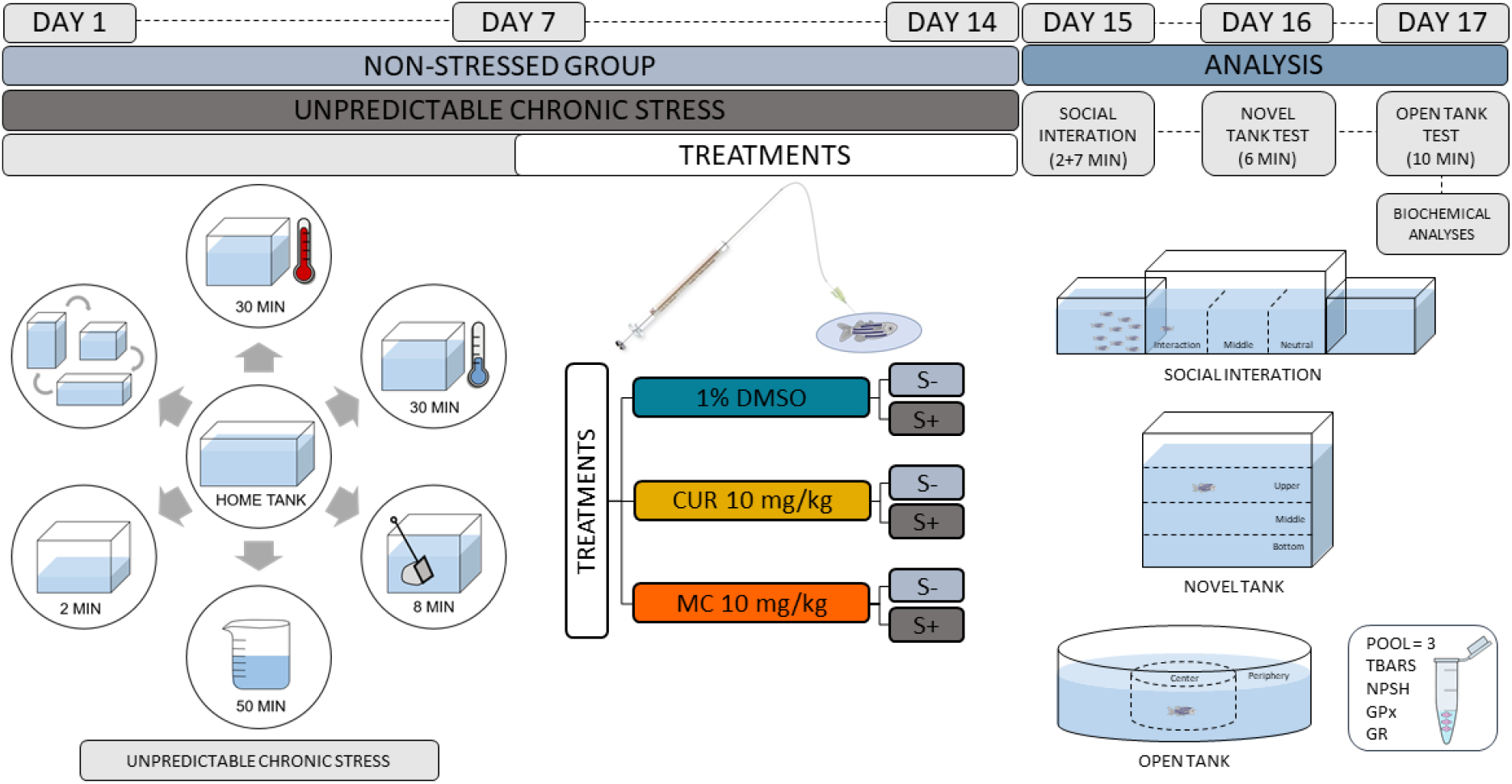
Experimental design. Zebrafish remained in the home tank and were subjected to the UCS for 14 days or remained undisturbed (non-stressed control). After the first seven days of stress, zebrafish were daily injected i.p. with DMSO 1%, CUR 10 mg/kg, or MC 10 mg/kg. On the 15^th^ of the experimental protocol, animals were subjected to the social interaction test. On the 16^th^ animals were subjected to the novel tank test. On the 17^th^ animals were subjected to the open tank test and then euthanized to collect the brain, which was used in biochemical analyses. DMSO (dimethyl sulfoxide), CUR (curcumin) and MC (micronized curcumin).

#### 2.8.1 Social interaction test (SI)

The SI test was conducted as described previously (Benvenutti et al., 2020; Bertelli et al., 2021) Animals were placed individually for 7 min in a tank (30 x 10 x 15 cm, 10 cm water level) flanked by two identical tanks (15 x 10 x 13 cm, 10 cm water level), either empty (neutral stimulus) or containing 10 unfamiliar zebrafish (social stimulus) (Fig. 3). The position of the social stimulus (right or left) was counterbalanced throughout the tests. The test apparatus was virtually divided into three vertical areas (interaction, middle, and neutral). Videos were recorded from the front view. Animals were habituated to the apparatus for 2 min and then analyzed for 5 min. The following parameters were quantified: total distance traveled, number of crossings (transitions between the areas of the tank), time spent in the interaction area (as a proxy for social interaction time), and time spent in the neutral area (Seibt et al., 2011). We changed the water in the tanks between animals to avoid interference from drug traces or alarm substances released by previously tested fish.

#### 2.8.2 Novel tank test (NTT)

The NTT was conducted as described previously (Benvenutti et al., 2020; Bertelli et al., 2021; Marcon et al., 2019; Mocelin et al., 2019). Animals were individually placed in the tank (24 × 8 × 20 cm, 15 cm water level) and recorded for 6 min. Videos were recorded from the front view. The test apparatus was virtually divided into three horizontal areas (top, middle, and bottom) (Marcon et al., 2019; Mocelin et al., 2019). The water in the tanks was changed between animals to avoid interference from drug traces or alarm substances released by previously tested fish. The following parameters were quantified: total distance traveled (m), number of crossings (transitions between the areas of the tank), time spent (s), and number of entries in the top area of the tank.

#### 2.8.3 Open tank test (OTT)

The OTT was conducted as described previously (Benvenutti et al., 2020; Bertelli et al., 2021). Animals were individually placed in the center of a circular arena made of opaque white plastic (24 cm diameter, 8 cm walls, 2 cm water level) and recorded for 10 min. The apparatus was virtually divided into two areas for video analyses: the central area of 12 cm in diameter and the periphery. Videos were recorded from the top view. The following parameters were quantified: total distance traveled (m), number of crossings (transitions between the areas of the tank), absolute turn angle (°), and time spent in the center area of the tank (s) (Johnson and Hamilton, 2017; Krook et al., 2019)

### 2.9 Biochemical assays

For each independent sample, three brains were pooled (n=6) and homogenized in 450 μL of phosphate-buffered saline (PBS, pH 7.4, Sigma-Aldrich) and centrifuged at 10,000 g at 4 °C in a cooling centrifuge; the supernatants were collected and kept in microtubes on ice until the assays were performed. The detailed protocol for prepare brain tissue samples is available at protocols.io(Sachett et al., 2020a). The protein was quantified according to the Coomassie blue method using bovine serum albumin (Sigma-Aldrich) as a standard (Bradford, 1976). The detailed protocol for protein quantification is available at protocols.io (Sachett et al., 2020b)

#### 2.9.1 Non-protein thiols (NPSH)

The content of NPSH in the samples was determined by mixing equal volumes of the brain tissue preparation (50 μg of proteins) and trichloroacetic acid (TCA, 6%), centrifuging the mix (10,000 g, 10 min at 4 °C), the supernatants were added to TFK (1 M) and DTNB (10 mM) and the absorbance was measured at 412 nm after 1 h. The detailed protocol is available at protocols.io (Sachett et al., 2020c).

#### 2.9.2 Glutathione reductase activity (GR)

The GR activity in the samples was determined by mixing the sample (30 μg of protein) with a reaction medium containing TFK + EDTA (154 mM, pH 7.0) and nicotinamide adenine dinucleotide phosphate (NADPH, 2mM). Then, oxidized glutathione (GSSG, 20 mM) was added and the increase of NADPH absorbance per minute was read at 340 nm. The detailed protocol is available at protocols.io (Sachett et al., 2021e)

#### 2.9.3 Glutathione peroxidase activity (GPx)

The GPx activity in the samples was determined by mixing the sample (30 μg of protein) with a reaction medium containing TFK + EDTA (0.5 M, pH 7.0), NADPH (1.6 mM), GSH (10 mM), GR (2.5 U/mL), and 10 mM azide. Then, H_2_O_2_ (4 mM) was added and the decrease of NADPH absorbance per minute was read at 340 nm. The detailed protocol is available at protocols.io (Sachett et al., 2021f).

#### 2.9.4 Substances reactive to thiobarbituric acid (TBARS)

The lipid peroxidation was evaluated by quantifying the production of TBARS. Samples (50 μg of proteins) were mixed with TBA (0.5%) and TCA (20%) (150 μL). The mixture was heated at 100 °C for 30 min. The absorbance of the samples was determined at 532 nm in a microplate reader. MDA (2 mM) was used as the standard. The detailed protocol is available at protocols.io (Sachett et al., 2020d).

### 2.10 Statistical analysis

We calculated the sample size to detect an effect size of 0.35 for the interaction between stress and treatment with a power of 0.9 and an alpha of 0.05 using G*Power 3.1.9.7 for Windows. The total distance traveled was defined as the primary outcome. The total sample size was 107, which was rounded up to 108 to yield n = 18 animals per experimental group.

The normality and homogeneity of variances were confirmed for all data sets using D’Agostino-Pearson and Levene tests, respectively. Results were analyzed by one-way (analysis of the antioxidant activity *in vitro*), or two-way (UCS) ANOVA followed by Tukey post hoc test when applicable. The outliers were defined using the ROUT statistical test and were removed from the analyses. This resulted in 6 outliers (1 animal from each group) removed from the SI test, 4 outliers (1 animal from CUR S-, MC S-, DMSO S+ and MC S+ groups) removed from the NTT and 2 outliers (1 animal from each DMSO S+ and MC S+ groups) removed from the OTT. The tank and sex effects were tested in all comparisons and no effect was observed, so the data were pooled.

Data are expressed as mean ± standard deviations of the mean (S.D.). The level of significance was set at p<0.05. Data were analyzed using IBM SPSS Statistics version 27 for Windows and the graphs were plotted using GraphPad Prism version 8.0.1 for Windows.

## 3. Results and discussion

### 3.1. Micronization

Characterization and size of the particles of the non-micronized and micronized curcumin are presented in Figure 4. The SEM showed a decrease in the size and change in the morphology of the CUR particles after micronization (Fig. 4A and 4B), being observed by the difference in zoom capable of visualizing the shape of the particles. CUR has an average size of 12.36 μm while the average size obtained through SEDS was 2.29 μm, which means a reduction of 5.4 times (Fig. 4C). The DSC data (Fig. 4D) showed the detected melting point for curcumin was 176.01 °C, while for micronized curcumin the melting point was reduced to 170.9 °C. Changes to the melting point are related to alterations in dissolution and solubility rates, among other properties, and are caused by modification in the crystalline structure of the composts (Aguiar et al., 2018, 2017, 2016; Bertoncello et al., 2018; Chen et al., 2012; Cheng et al., 2016; Li et al., 2015; Moribe et al., 2005; Zhang et al., 2009).

**Fig. 4.**
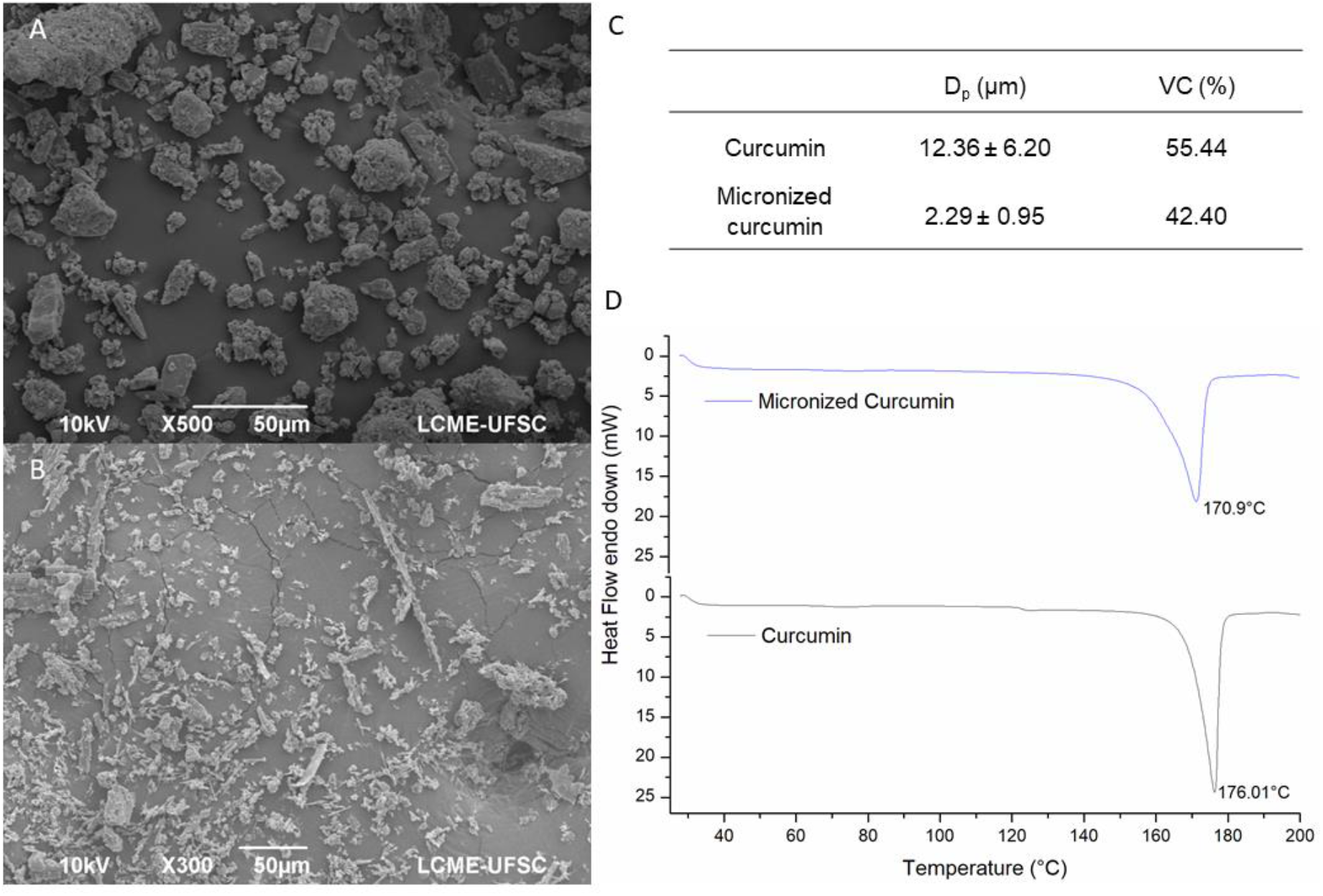
Effects of micronization of curcumin on size values and thermal analysis. (A) particle size in the scanning electron microscope of non-micronized curcumin, (B) particle size in the scanning electron microscope of micronized curcumin, (C) particle size values in raw compounds, (D) differential scanning calorimetry. Dp (average particle diameter); VC (variation coefficient).

The micronization of curcumin (Bertoncello et al., 2018), N-acetylcysteine (Aguiar et al., 2017), trans-resveratrol (Aguiar et al., 2018, 2016; Almeida et al., 2021; Decui et al., 2020), methotrexate (Chen et al., 2012), phenylbutazone (Moribe et al., 2005), *Panax notoginseng* saponins (Liang et al., 2021), etoposide (Cheng et al., 2016), ellagic acid (Li et al., 2015), taxifolin (Zu et al., 2012) atorvastatin calcium (Zhang et al., 2009) and ibuprofen (Han et al., 2011; Sosna et al., 2018) by the SEDS technique have shown a reduction in particle size, an increase in the dissolution rate, an increase in the solubility, as well as a modification of the crystalline structure of the compound when compared with non-micronized composts. Recently, the micronization of *Panax notoginseng* saponins changed the pharmacokinetic parameters of these compounds in rats, showing significantly higher values in plasma powders with smaller particle sizes than the larger particle sizes (Liang et al., 2021). Therefore, micronization can improve the bioavailability of compounds, being a critical tool for the industry, especially for compounds with low solubility (Aguiar et al., 2016; Chau et al., 2007; Chen et al., 2012; Li et al., 2015).

### 3.2. Antioxidant activity in vitro

The experimental design is summarized in Figure 2. In this experiment, we analyzed the *in vitro* antioxidant effects of CUR and MC in comparison to the positive control ascorbic acid. The FRAP method quantifies the electron-donating capacity of a compound by reduction of iron from ferric status to ferrous in solution (Halliwell, 1990). MC (at 0.25 and 1 g/L) increased the reduction of iron compared to CUR in the same concentrations (p <0.0001, F_9,40_ = 316.2, Fig. 5A). As expected, ascorbic acid showed a greater reducing capacity than MC and CUR in all concentrations. MC and ascorbic acid in all concentrations and CUR at 0.25 and 1 g/L exhibited an increased reduction potential when compared to 1% DMSO. These results indicate that the electron-donating capacity was increased by micronization. Similar results have been found in the literature where micronization increased the iron-reducing capacity (Li et al., 2015; Lu et al., 2020; Zhu et al., 2014).

**Fig. 5.**
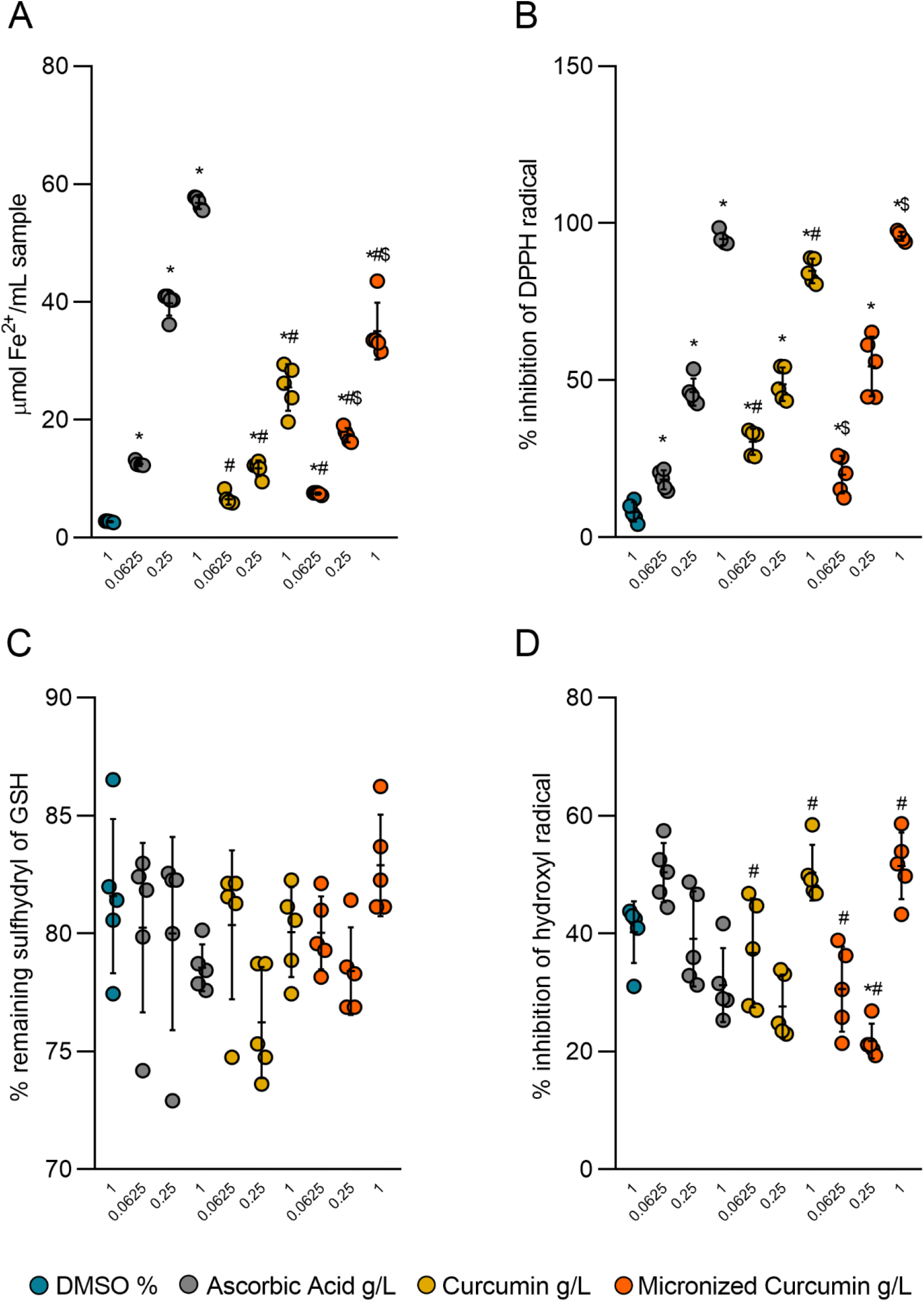
Effects of DMSO, Ascorbic Acid, Curcumin, and Micronized curcumin on antioxidant activity *in vitro*. Data are expressed as mean ± S.D. One-way ANOVA/Tukey. n= 5. ^*^p<0.05 x 1% DMSO. ^#^p<0.05 x ascorbic acid (in the same concentration). ^$^p<0.05 x curcumin (in the same concentration). DMSO (dimethyl sulfoxide), FRAP (ferric reducing antioxidant power), DPPH (1,1-diphenyl-2-2-picyryl-hydrazyl), and GSH (L-Glutathione reduced).

DPPH is a free radical used to evaluate the antioxidant capacity through its elimination by reduction via donation of a hydrogen atom by a compound (Brand-Williams et al., 1995). In the DPPH assay, MC 1 g/L increased the scavenging of the DPPH radical when compared to CUR at the same concentration, being equivalent to the positive control ascorbic acid at all concentrations tested (p<0.0001, F_9,40_ = 228.2, Fig. 5B). CUR 0.0625 g/L removed more radicals than MC and ascorbic acid at the same concentration, but less than ascorbic acid at 1 g/L. MC, CUR, and ascorbic acid, in all concentrations tested, showed more inhibition of the radical than DMSO. There was no statistical difference in the EC_50_ among the treatments. Other studies reported that resveratrol, NAC, taxifolin, ellagic acid, and apple pomace micronized by SEDS, increased the antioxidant activity when compared to the non-micronized compost in DPPH tests (Aguiar et al., 2018, 2017; Li et al., 2015; Lu et al., 2020; Zu et al., 2012).

There were no significant differences between treatments on the percentage of remaining sulfhydryl groups of GSH (Fig. 5C). However, all treatments were able to inhibit oxidation of GSH measured by the remaining sulfhydryl groups of GSH that react with DTNB after oxidation induced by H_2_O_2_.

In the deoxyribose assay, OH· generated by Fenton reaction, oxidizes deoxyribose with the formation of MDA, quantified by TBARS (Halliwell et al., 1987). Thus, the ability of the compound to inhibit the formation of OH· and, consequently, the inhibition of the formation of MDA was analyzed. Both MC and CUR 1 g/L increased the inhibition of OH· production when compared to ascorbic acid (p <0.0001, F_9,40_ = 13.92, Fig. 5D). However, ascorbic acid 0.0625 g/L showed a greater inhibition when compared to CUR and MC at the same concentration and, also ascorbic at 2.5 g/L when compared to MC 2.5 g/L. MC 0.25 g/L showed less inhibition than 1% DMSO.

These results indicate that curcumin can eliminate peroxides, inhibit the formation of free radicals, and protect against an imbalance in the oxidative state of the organism and lipid peroxidation. Studies indicate that the antioxidant effect of curcumin can be attributed to its phenolic groups, acting as reducing agents, hydrogen donors, as well as oxygen adsorption inhibitors (Zheng et al., 2017). This supports our results suggesting that the ability to eliminate free radicals from MC obtained by the DPPH and deoxyribose assay is related to their reducing properties in the FRAP. Similar results have already been observed in several studies using curcumin preparations *in vitro* (Landeros et al., 2017; Sahu, 2016; Singh et al., 2018). However, this is the first study evaluating these effects in micronized curcumin.

Studies using this technology showed that the SEDS process is a robust methodology for improving the physicochemical properties and antioxidant activity by increasing the bioaccessibility and availability of phenolic and other compounds active (Lu et al., 2020; Sefrin Speroni et al., 2021; Zhou et al., 2004; Zu et al., 2012). Furthermore, positive correlations were detected between radical scavenging activity, ferric reducing antioxidant power, and total phenolic content of wine grape pomace micronized (Zhu et al., 2014). These data suggest the possibility of improving the antioxidant effects of composts by micronization.

### 3.3 Effects of CUR and MC on behavioral and neurochemical parameters in zebrafish submitted to unpredictable chronic stress (UCS)

The timeline and experimental design are shown in Figure 3. Based on the most effective concentration of curcumin (1 g/L) observed in the *in vitro* antioxidant activity assays, we performed the UCS to verify the effects of both preparations on behavioral and neurochemical parameters in zebrafish. The unpredictable chronic stress induces several behavioral and neurochemical characteristics that resemble those observed in patients with anxiety and/or mood disorders (Chattarji et al., 2015; Willner, 2017, 2005). Antidepressants (Demin et al., 2020; Marcon et al., 2016; Reddy et al., 2021; Song et al., 2018), anxiolytics (Marcon et al., 2016), antioxidant (Marcon et al., 2019; Mocelin et al., 2019), ketamine (Reddy et al., 2021) and prazosin (O’Daniel and Petrunich-Rutherford, 2020) showed protective effects in this model.

Zebrafish live in social groups and, like humans, respond to the social support of conspecifics from a familiar shoal and recover from stressful events better in the presence of conspecifics (Faustino et al., 2017). In the SI test, two-way ANOVA revealed a main effect of stress on total distance (Fig. 6A), indicating a locomotor dysfunction induced by UCS. Both CUR and MC were unable to block this effect. The number of crossings (Fig. 6B), time in the interaction (Fig. 6C), and neutral (Fig. 6D) areas were not affected by UCS, indicating that, in this experimental condition, there was no impact of stress on social preference parameters. Notably, in the scientific literature, the effect of UCS on zebrafish shoaling behavior has been inconsistent, presenting decreased or increased social interaction after UCS (Chakravarty et al., 2013; Piato et al., 2011). In contrast to these studies, Fulcher et al. (2017) and Bertelli et al., (2021) found that UCS did not alter the social interaction of zebrafish in the social interaction test.

**Fig. 6.**
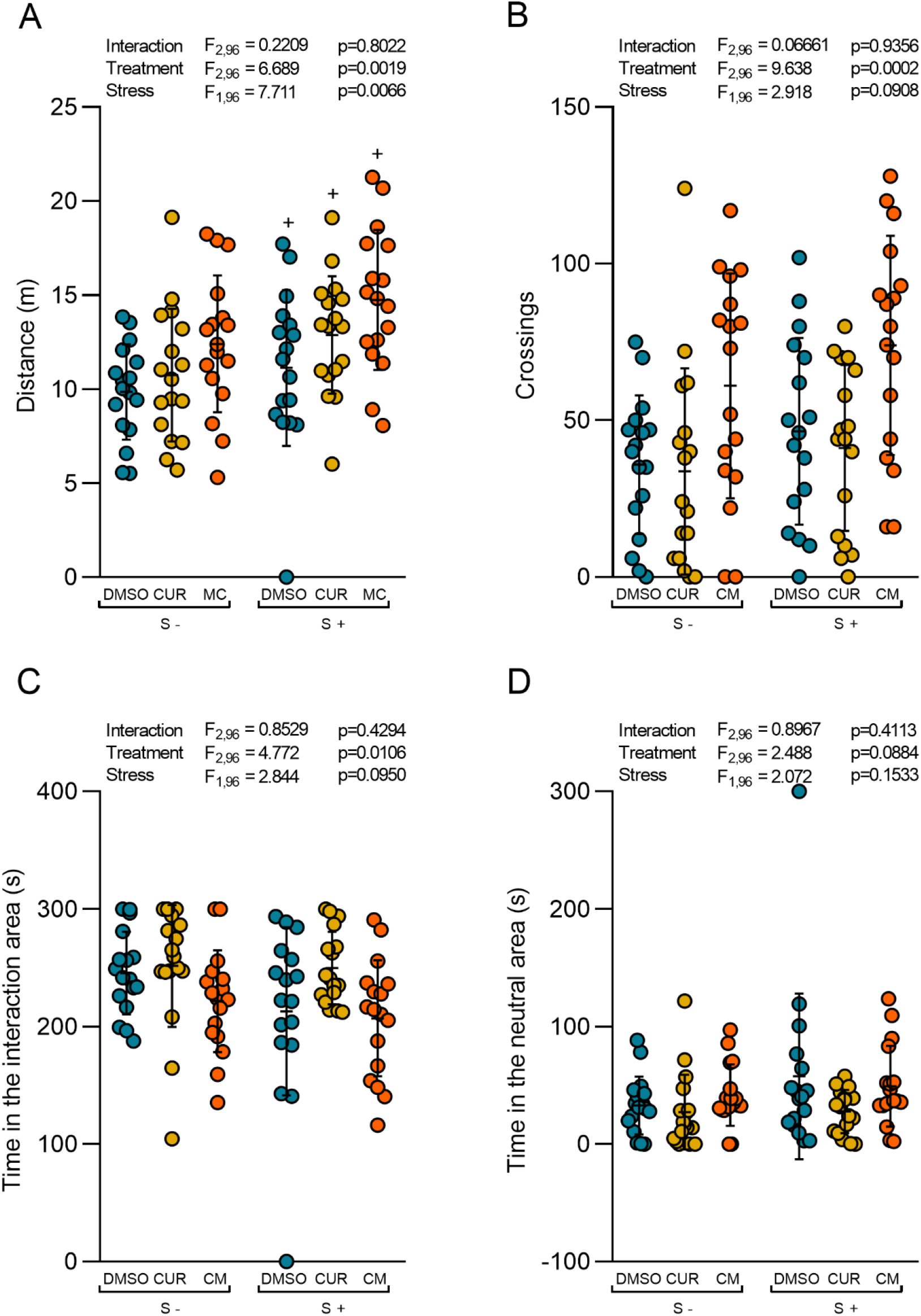
Effects of CUR and MC (10 mg/kg) in zebrafish submitted to the UCS on the social interaction test (day 15). (A) total distance traveled (B) the number of crossings, (C) time in the interaction area and (D) time in the neutral area. Data are expressed as mean ± S.D. Two-way ANOVA/Tukey. n=17 ^+^p<0.05 stress effect. DMSO (dimethyl sulfoxide); CUR (curcumin); MC (micronized curcumin).

In the NTT, a reduction of the exploratory behavior towards the top area can be interpreted as an index of anxiety. Anxiolytic drugs such as fluoxetine, buspirone, and diazepam increase, whereas anxiogenic drugs such as caffeine and nicotine decrease the time on the top area of the tank (Bencan et al., 2009; Egan et al., 2009; Gebauer et al., 2011; Levin et al., 2007). Here, UCS decreased the number of crossings (Fig. 7B), time spent (Fig. 7C), and entries (Fig. 7D) to the top area of the tank in the NTT. Neither the UCS nor the treatments altered the total distance traveled (Fig. 7A). These results replicate and reinforce previous studies showing that UCS induces anxiety-like behavior in zebrafish (Chakravarty et al., 2013; Demin et al., 2020; Marcon et al., 2019, 2018a, 2016; Mocelin et al., 2019; O’Daniel and Petrunich-Rutherford, 2020; Piato et al., 2011; Reddy et al., 2021; Song et al., 2018). Both CUR and MC were unable to block these alterations.

**Fig. 7.**
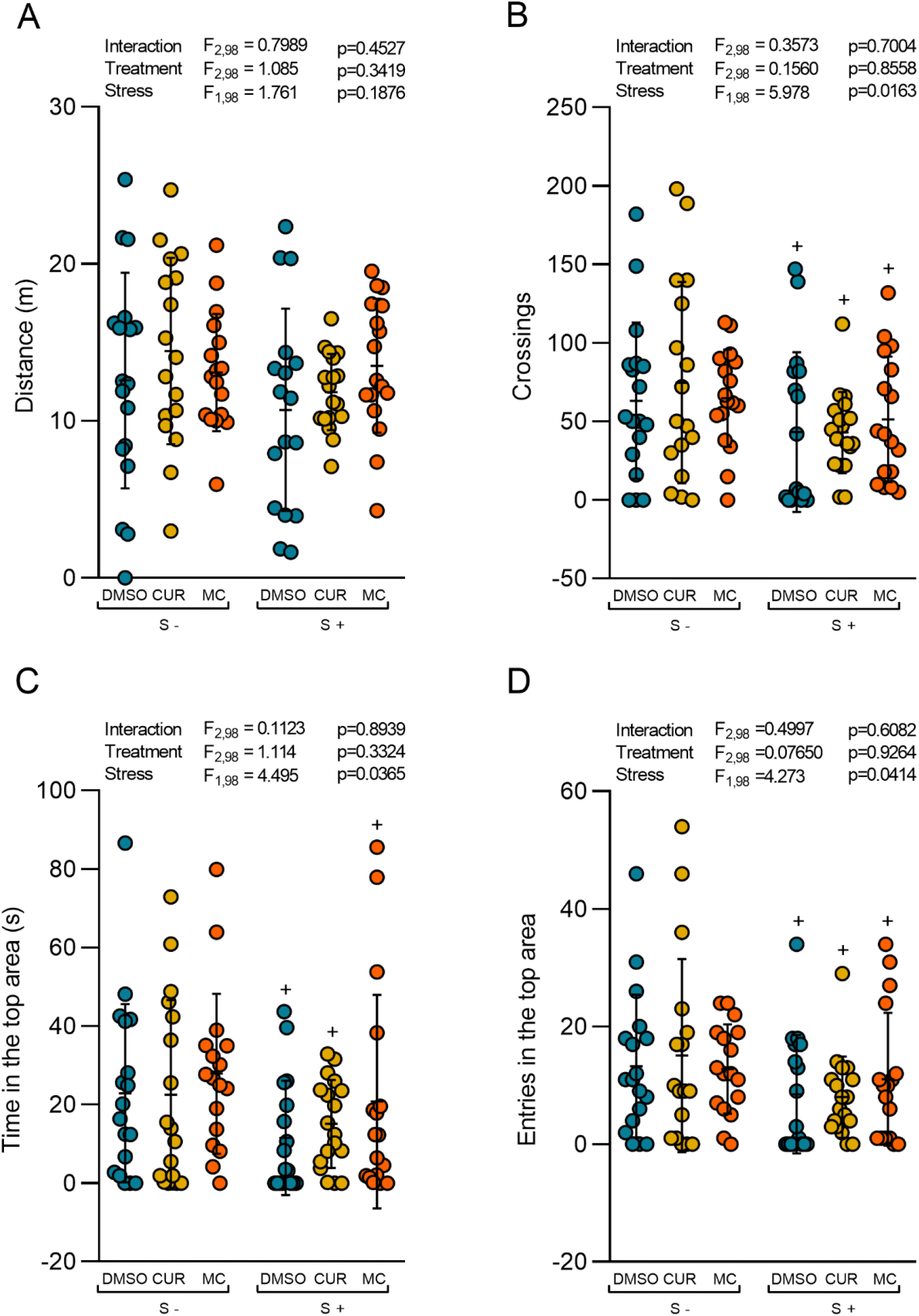
Effects of CUR and MC (10 mg/kg) in zebrafish submitted to the UCS on the novel tank test (day 16). (A) total distance traveled, (B) the number of crossings, (C) time in the top area, (D) entries in the top area. Data are expressed as mean ± S.D. Two-way ANOVA/Tukey. n=17-18. ^+^p<0.05 stress effect. DMSO (dimethyl sulfoxide); CUR (curcumin); MC (micronized curcumin).

The OTT is a paradigm adapted from the open-field test (OFT) used in rodents, showing similarity with exploration, thigmotaxis, and freezing parameters. A decrease in the time spent in the thigmotaxis area (periphery) and an increase in exploration indicates a decrease in anxiety in zebrafish (Johnson and Hamilton, 2017; Stewart et al., 2012). Two-way ANOVA revealed an interaction between both factors on the number of line crossings (Fig. 8B), however, no significant effects were observed in *post hoc* analysis. In addition, there were no significant effects of any intervention on total distance, absolute turn angle, and time spent in the center area (Fig. 8A, 8C, and 8D, respectively).

**Fig. 8.**
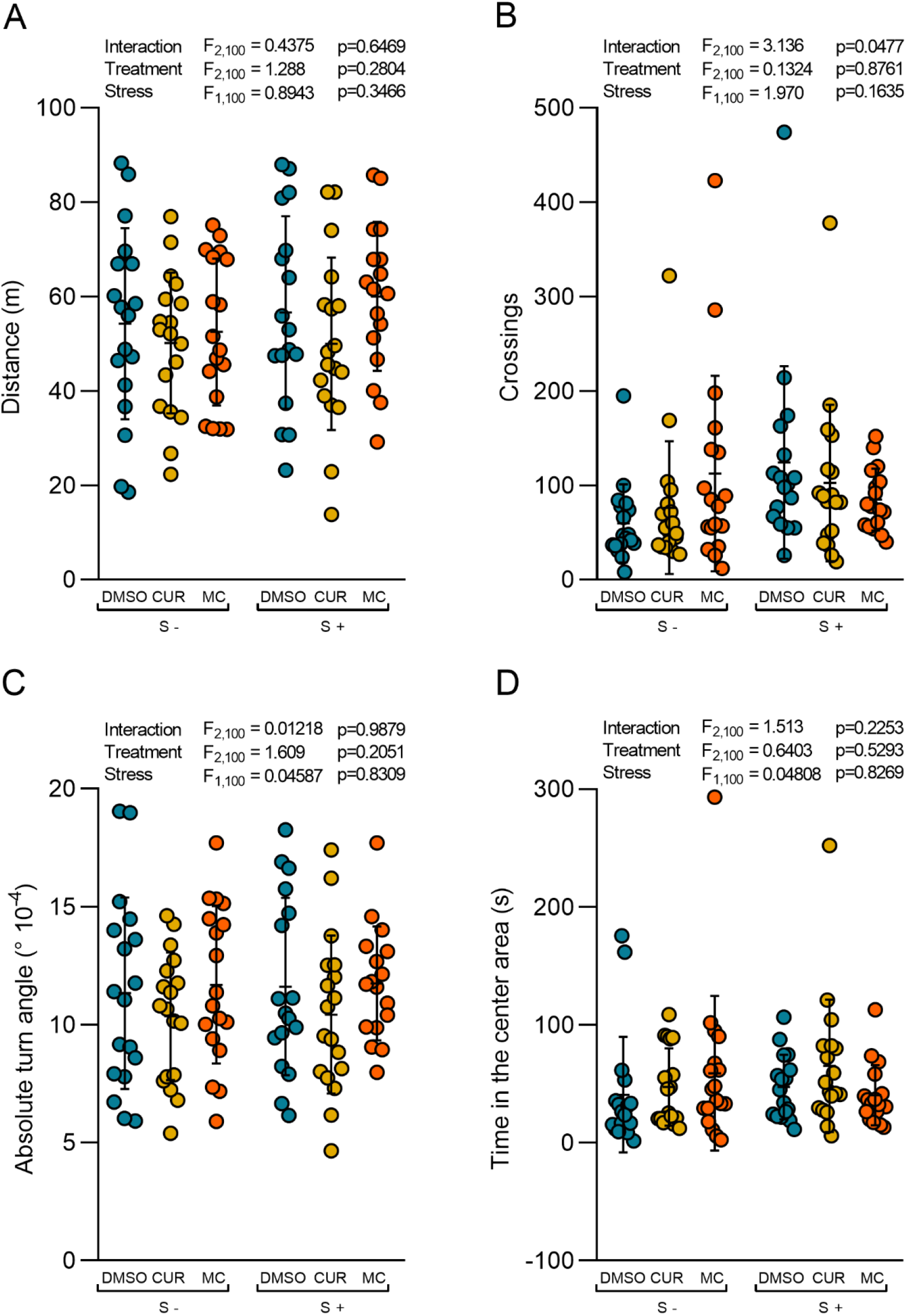
Effects of CUR and MC (10 mg/kg) in zebrafish submitted to the UCS on the open tank test (day 17). (A) total distance traveled (B) the number of crossings, (C) absolute turn angle, (D) time in the center area. Data are expressed as mean ± S.D. Two-way ANOVA. n=17-18. DMSO (dimethyl sulfoxide); CUR (curcumin); MC (micronized curcumin).

Several studies evaluated the effects of curcumin using animal models of chronic stress. In rodents submitted to stress for 21 or 28 days, curcumin (10-40 mg/kg, i.p. or p.o. for the same time) blocked the stress effects decreasing immobility time in the OFT and memory deficits in the Morris water maze (MWM). In these studies, curcumin decreased serum corticosterone and increased BDNF and monoamine levels. Moreover, curcumin increased hippocampal neurogenesis and decreased brain monoamine oxidase activity, when compared to the stressed group (Bhutani et al., 2009; da Silva Marques et al., 2021; Xu et al., 2009, 2007, 2006). In rats submitted to UCS for 35 days, curcumin (40 mg/kg i.p. for 35 days) decreased the immobility time and increased the swimming time in the forced swim test, besides increased the percent of sucrose consumption (Fan et al., 2019, 2018). Similarly, treatment with curcumin (20 mg/kg p.o.) for 8 days increased the glucose preference and locomotor activity in OFT, as well as a decrease in the escape latency in MWM and the brain cytokines levels in rats submitted to UCS for 8 days (Vasileva et al., 2018). We supposed that the effects of curcumin on stress-induced behavioral changes might be observed with a longer exposure time or even in a dose range different from that used in the present study.

After the behavioral tests, we evaluated the effects of CUR and MC on neurochemical parameters in zebrafish submitted to UCS. Two-way ANOVA revealed an interaction between both factors for NPSH levels, GPx and GR enzyme activity, and TBARS levels. The post hoc analysis showed that UCS decreased the NPSH levels (a measure that reflects the levels of GSH) (Fig. 9A), increased the GR enzyme activity (Fig. 9B), and increased lipid peroxidation (measured by TBARS levels) (Fig. 9D). These results indicate an oxidative status disturbance UCS-induced and consequently oxidative damage in the zebrafish brain. Despite the two-way ANOVA revealed an interaction between both factors on the GPx enzyme activity, no significant effects were observed in post hoc analysis between the control non-stressed and stressed group. MC blocked the effects of UCS, normalizing GR activity, and increasing NPSH levels and GPx activity, consequently decreasing the lipid peroxidation. CUR was able to increase GPx enzyme activity, although it was unable to block the effects of the stress on lipid peroxidation, NPSH levels, and GR activity.

**Fig. 9.**
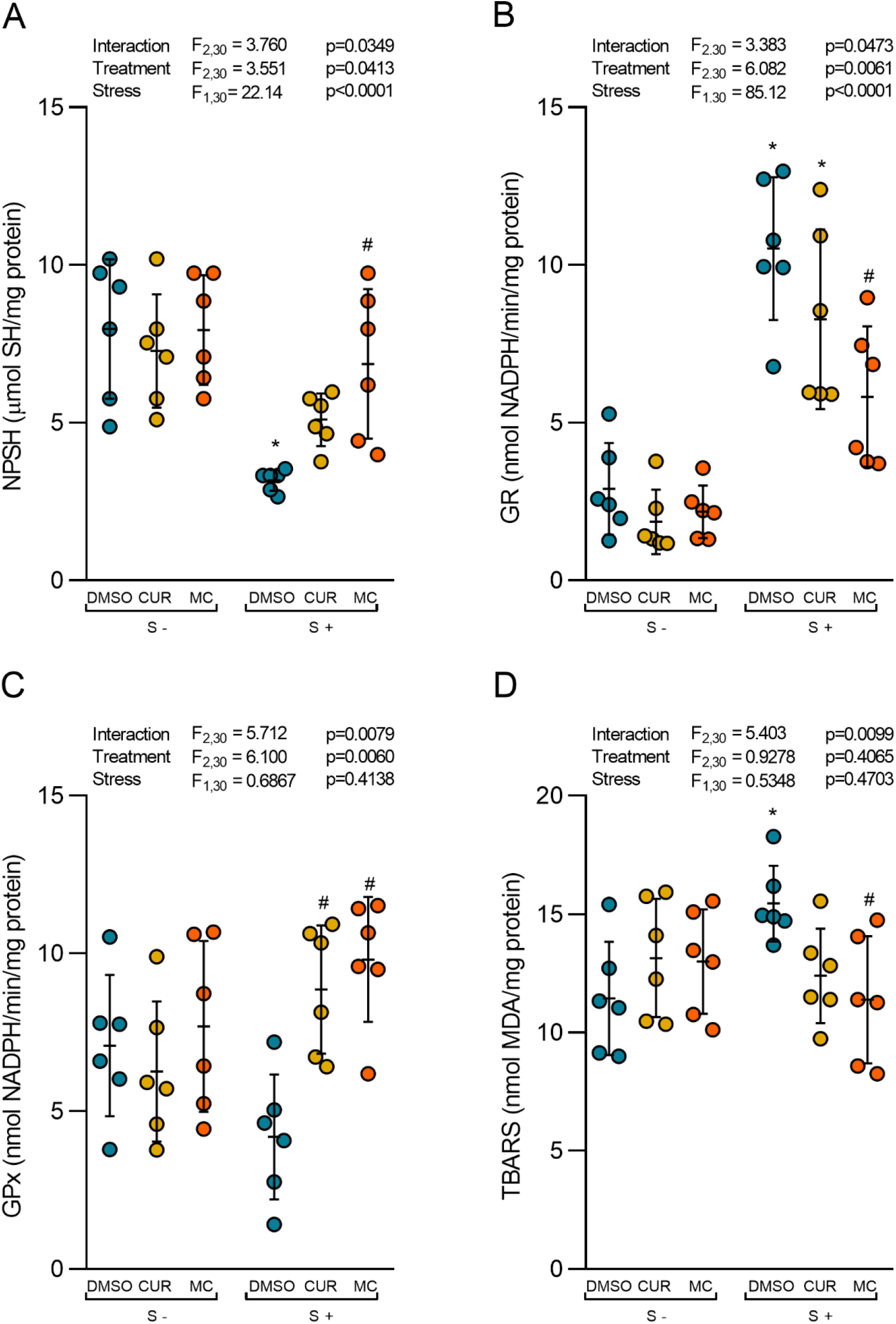
Effects of CUR and MC (10 mg/kg) in zebrafish submitted to UCS on neurochemical parameters. (A) non-protein thiol levels, (B) glutathione reductase activity, (C) glutathione peroxidase activity, (D) lipid peroxidation levels. Data are expressed as mean ± S.D. Two-way ANOVA/Tukey. n=6. ^*^p<0.05 x 1% DMSO non-stressed. ^#^p<0.05 x 1% DMSO stressed. DMSO (dimethyl sulfoxide); CUR (curcumin); MC (micronized curcumin).

Some organs, like the brain, are more vulnerable to the detrimental effects of ROS because it has a high metabolic rate and lower antioxidant levels (Maes et al., 2011; Mandelker, 2008). Stress adaptation or allostasis increases cerebral energy demand, reflecting the increased mitochondrial activity within the brain, which is related to high oxygen consumption and greater ROS production (Avery, 2011; Harwell, 2007; Picard et al., 2018). Moreover, the increased production of ROS has been linked to hyperactivation of the HPA axis with a consequent increase in cortisol secretion by producing damage to hippocampal neurons, which maintain the homeostasis of the HPA axis by negative feedback mechanisms (Bhatia et al., 2011; KVETŇANSKÝ et al., 1995; Maes et al., 2011; SAPOLSKY et al., 1986). The overproduction of ROS and decrease in antioxidant defenses consequently cause an oxidative stress status that alters neuronal homeostasis and favors the occurrence of oxidative lesions in proteins, lipids, and nucleic acids, which may result in cell death (Avery, 2011; Harwell, 2007; Valko et al., 2007). The reduced GSH is one of the main antioxidant components involved in the removal of ROS and maintenance of oxidative status. GPx reduces H_2_O_2_ through the GSH oxidation to oxidized glutathione in dimerized form (GSSG). GSSG is then recycled by the enzyme GR through the NADPH oxidation reaction to oxidized NADP. In addition, GSH also can undergo oxidation and form disulfides of the GSSR type with the cysteine thiol present in proteins. The sulfhydryl group present in the cysteine thiol is the active site and is responsible for its protective functions against oxidative stress. Therefore, its oxidation leads to the formation of GSH disulfides and inactivation of its antioxidant capacity, leaving the organism more susceptible to suffer oxidative damage (Dasuri et al., 2013; Gandhi and Abramov, 2012).

Our results suggest that unpredictable chronic stress decreased the antioxidant defenses by depleting cerebral GSH, making the zebrafish brain more susceptible to oxidative damage such as lipid peroxidation. On the other hand, UCS increased the activity of GR as a reflex of the organism in face of the decrease in GSH. Indeed, studies have shown the UCS increases body cortisol levels, at the same time that increase ROS production, decreases the antioxidant mechanisms (NPSH level and antioxidant enzyme activity superoxide dismutase (SOD)), and consequently increases lipid peroxidation (TBARS levels) in the zebrafish brain (Marcon et al., 2019, 2018b, 2018a; Mocelin et al., 2019). Interestingly, we also demonstrated for the first time that MC has a better protective effect than CUR against oxidative stress, blocking the stress-induced neurochemical effects in the zebrafish brain. We suggest that CM prevented oxidative stress by increasing antioxidant defenses (GSH level and GPx enzyme activity).

## 4. Conclusion

In this study, we have shown micronized curcumin prevented the effects of chronic stress on neurochemical markers, despite the absence of effects on behavioral parameters. Considering the heterogeneous and complex effects caused by chronic stress, one possibility is that curcumin would be exerting neuroprotective effects as an antioxidant, but not being able to modulate other systems involved in stress-induced behavioral changes. However, there is a clear superiority of the micronized preparation over the conventional against stress-induced oxidative stress. Finally, the micronization with the SEDS technique altered the crystalline structure of the compound and its melting point, which led to significant improvements in both *in vivo* and *in vitro* tests. Thus, the micronization may increase bioavailability and potentiate the therapeutic effect of drugs, making the technology of supercritical fluid micronization promising to the pharmaceutical industry.

## Supporting information

Supplemental Table S1 Sachett et al

