## Supplemental Table S1 Sachett et al for "Curcumin micronization by supercritical fluid: *in vitro* and *in vivo* biological relevance"

### Supplementary Material

#### Micronized curcumin blocks the effects of unpredictable chronic stress on neurochemical but not behavioral outcomes in zebrafish

Adrieli Sachett[A], Matheus Gallas-Lopes[B], Radharani Benvenutti[A], Matheus Marcon[A], Gean Pablo S. Aguiar[D], Ana Paula Herrmann[B,C], J. Vladimir Oliveira[D,E], Anna M. Siebel[D], Angelo Piato[A,B]*

[A] Programa de Pós-graduação em Neurociências, Instituto de Ciências Básicas da Saúde, Universidade Federal do Rio Grande do Sul (UFRGS), Porto Alegre, RS, Brazil.
[B] Departamento de Farmacologia, Instituto de Ciências Básicas da Saúde, Universidade Federal do Rio Grande do Sul (UFRGS), Porto Alegre, RS, Brazil.
[C] Programa de Pós-graduação em Farmacologia e Terapêutica, Instituto de Ciências Básicas da Saúde, Universidade Federal do Rio Grande do Sul (UFRGS), Porto Alegre, RS, Brazil.
[D] Programa de Pós-Graduação em Ciências Ambientais, Universidade Comunitária da Região de Chapecó (Unochapecó), Chapecó, SC, Brazil.
[E] Departamento de Engenharia Química e de Alimentos, Universidade Federal de Santa Catarina (UFSC), Florianópolis, SC, Brazil.

*Correspondence to: Angelo Piato, Ph.D. Departamento de Farmacologia, Instituto de Ciências Básicas da Saúde, Universidade Federal do Rio Grande do Sul (UFRGS), Av. Sarmento Leite, 500/305, Porto Alegre, RS, 90050-170, Brazil; Phone/Fax: +55 51 33083121; E-mail address:

**Table S1.** Schedule and stressors of the unpredictable chronic stress protocol.

| | Day/Time | 1 | 2 | 3 | 4 | 5 | 6 | 7 |
|---|---|---|---|---|---|---|---|---|
| STRESS | Morning | **10:30 AM** Tank change (3 times / 10 min each) | **08:00 AM** Overcrowding (9 animals in 200 mL beaker / 50 min) | **08:30 AM** Low housing tank water level until dorse exposure (2 min) | **09:30 AM** Heating tank water (33 ºC / 30 min) | **08:30 AM** Cooling tank water (23 ºC / 30 min) | **09:15 AM** Chasing with a net (8 min) | **8:00 AM** Tank change (3 times / 10 min each) |
| | Afternoon | **03:00 PM** Chasing with a net (8 min) | **04:30 PM** Low housing tank water level until dorse exposure (2 min) | **04:00 PM** Cooling tank water (23 ºC / 30 min) | **03:30 PM** Overcrowding (9 animals in 200 mL beaker / 50 min) | **04:00 PM** Tank change (3 times / 10 min each) | **04:30 PM** Low housing tank water level until dorse exposure (2 min) | **03:30 PM** Heating tank water (33 ºC / 30 min) |

| | Day/Time | 8 | 9 | 10 | 11 | 12 | 13 | 14 |
|---|---|---|---|---|---|---|---|---|
| STRESS + TREATMENT | Morning | **09:15 AM** Chasing with a net (8 min) | **08:00 AM** Tank change (3 times / 10 min each) | **09:15 AM** Cooling tank water (23 ºC / 30 min) | **08:30 AM** Overcrowding (9 animals in 200 mL beaker / 50 min) | **09:15 AM** Low housing tank water level until dorse exposure (2 min) | **08:15 AM** Heating tank water (33 ºC / 30 min) | **10:00 AM** Overcrowding (9 animals in 200 mL beaker / 50 min) |
| | Afternoon | **04:00 PM** Overcrowding (9 animals in 200 mL beaker / 50 min) | **03:00 PM** Chasing with a net (8 min) | **05:00 PM** Heating tank water (33 ºC / 30 min) | **03:30 PM** Chasing with a net (8 min) | **04:30 PM** Cooling tank water (23 ºC / 30 min) | **04:45 PM** Low housing tank water level until dorse exposure (2 min) | **04:00 PM** Tank change (3 times / 10 min each) |

| | Day/Time | 15 | 16 | 17 |
|---|---|---|---|---|
| STRESS + TREATMENT + TESTS | Morning | **08:00 AM** Social interaction test | **08:00 AM** Novel tank test | **08:00 AM** Open tank test and euthanasia |
| | Afternoon | **04:00 PM** Heating tank water (33 ºC / 30 min) | **04:30 PM** Cooling tank water (23 ºC / 30 min) | - |
